# Integrating eDNA and acoustic-trawl data to provide small pelagic biomass estimates for fisheries assessment

**DOI:** 10.1101/2025.06.11.659118

**Authors:** Cristina Claver, Beatriz Sobradillo, Iñaki Mendibil, Oriol Canals, Guillermo Boyra, Leire Ibaibarriaga, Ryan P. Kelly, Naiara Rodríguez-Ezpeleta

## Abstract

Accurate abundance estimates of fisheries resources are essential for sustainable fisheries management. In response to the growing need for developing more accurate and cost-effective biomass estimation methods, the analysis of environmental DNA (eDNA) has recently emerged as an alternative for fish abundance quantification. However, practical approaches for integrating eDNA data into fisheries assessment remain limited. Here, we introduce a Bayesian joint model that combines acoustic-trawl and eDNA data to estimate fish biomass. Utilizing 209 water eDNA samples and 196 acoustic transects, the model was applied to estimate the distribution and abundance of the European anchovy (*Engraulis encrasicolus*) in the Bay of Biscay. The joint model produced estimates consistent with known spatial patterns of anchovy, with eDNA data suggesting a broader distribution and potentially higher abundance. This research demonstrates the value of incorporating eDNA data as a complement to acoustic-trawl for stock assessment and illustrates the versatility of joint Bayesian models and their potential application to various species and datasets. Ultimately, our work opens new avenues for more holistic fisheries assessment, underscoring the growing role of eDNA in that context.

## Introduction

Pelagic fish play key ecological roles in marine ecosystems and constitute an important global food resource (Fréon *et al*. 2005), sustaining worldwide economically important fisheries (FAO 2024). Determining “how much fish there is” serves to sustainably manage these marine resources, safeguarding the long-term availability and proper ecosystem functioning. Abundance estimates are central to fisheries assessment, in particular for small pelagics (e.g., anchovies, sardines, herrings), which are characterized by largely fluctuating biomasses and highly dependent on environmental features (Checkley Jr *et al*. 2017). This makes them particularly vulnerable to overfishing and prone to collapse (Borja *et al*. 2019).

In last decades, trawl-acoustic methodology has become widespread to estimate abundance and spatial distribution of small pelagic fish globally (Barange *et al*. 2009). The method combines detection of acoustic signals measured continuously throughout the water column at high longitudinal and vertical image resolution (Simmonds and MacLennan 2008), with pelagic trawls providing data on species composition. Importantly, information about age and size of individuals can only be obtained through the catches derived from the trawling operations. Overall, the method has proven to be effective to cover large-scale areas (Fleischer 2005, Stenevik *et al*. 2015), but, at the same time, trawling has the disadvantages of being invasive, time consuming and dependant on catchability.

In recent years, the analysis of genetic material collected from marine water samples (environmental DNA or eDNA) has demonstrated to be suitable for assessing diversity, distribution and abundance of fish. This approach requires sampling a few litres of water, which makes it a low effort, non-invasive and cost-effective option, comparing to others. Beyond the benefits of easy sampling, eDNA has provided results consistent with those obtained from other established methods and it has outperformed them in some cases. For example, the analysis of eDNA collected from water samples through metabarcoding (i.e., the simultaneous taxonomic assignment of the DNA sequences obtained) results in species composition data that correlates with that obtained from catches, while detecting, in general, many more species (Thomsen *et al*. 2016, Stoeckle *et al*. 2021, Maiello *et al*. 2022, Veron *et al*. 2023). Also, relative abundances obtained through this approach coincide with those derived from the catches (Fraija-Fernández *et al*. 2020, Zhou *et al*. 2022). Consequently, eDNA metabarcoding has been promoted as a promising solution to improve fisheries research by increasing detectability while reducing invasiveness and costs (Lacoursière-Roussel *et al*. 2016).

Beyond qualitative detections and proportional data, there is a growing interest for the quantification of fish abundance through their eDNA (Gilbey *et al*. 2021) because the amount of eDNA has shown to consistently correlate with the number of fish and their biomass under controlled conditions (Takahara *et al*. 2012, Klymus *et al*. 2015, Eichmiller *et al*. 2016, Karlsson *et al*. 2022, Zhang *et al*. 2024). These studies rely on single species detection assays, which are more sensitive than metabarcoding and provide absolute and not only relative eDNA abundance data, which is more directly comparable with biomass. One notable and pioneering study is that of Shelton *et al*. (2022), which developed a biomass index modelling estimates of hake abundance based on eDNA concentration in a large oceanic area that were equivalent to acoustic estimates, demonstrating the capacity of eDNA to inform marine fishery assessment as efficiently as well-stablished data.

For now, most of the studies attempting to integrate eDNA abundance data in a fisheries assessment context are limited to comparing abundances derived from eDNA data with those obtained from other methods (Knudsen *et al*. 2019, Salter *et al*. 2019, Maes *et al*. 2024), and therefore little effort is focused to assess how to effectively integrate eDNA derived abundance estimates into the assessment. In this regard, recent work has shown that eDNA-based and traditional-based abundance data provide more comprehensive and accurate estimations when they are used together and some studies are now opting for combining rather than comparing methods (Guri *et al*. 2024, Chevrinais *et al*. 2025). Nonetheless, the quantitative combination of eDNA and acoustic-trawl data to estimate fish abundance has not been explored yet and it might be key to incorporate eDNA data into assessment of small pelagics.

In the North East Atlantic Ocean, the Bay of Biscay (BoB) is a temperate gulf where some of the most productive European small pelagic fisheries occur, including the anchovy (*Engraulis encrasicolus*) (ICES 2022), which underwent a closure (2005-2009) due to a stock collapse. Since 2003 acoustic surveys taking place in early autumn (JUVENA) are carried out yearly with the aim of monitoring the juveniles using trawl-acoustic methods (Boyra *et al*. 2008). After recovery, the anchovy fishery regulation in this area is based on Total Allowable Catch (TAC) established according to an international management plan agreed by different stakeholders (Sánchez *et al*. 2019, Uriarte *et al*. 2023). Being such an important and fragile resource, it is of special interest to explore new tools that could improve current monitoring methodologies. Here, we have used both acoustic-trawl and eDNA-derived information collected during the JUVENA surveys to develop a joint Bayesian statistical model to obtain spatial biomass estimates of European anchovy in the BoB, demonstrating the capacity of eDNA to provide valuable, additional information when combined with acoustic data.

## Materials and methods

### Acoustic-trawl sampling and data processing

Data analysed in this study was collected in the BoB over a period of 6 years (2018 to 2023) during the JUVENA acoustic surveys. This oceanographic campaign has taken place every year since 2003 in early autumn with the aim of estimating the abundance and spatial distribution of anchovy juveniles to quantify the annual recruitment (Boyra *et al*. 2013).

Details about the acoustic-trawl data processing methodology are explained in Boyra *et al*. (2023) and in Doray *et al*. (2021). Briefly, the strategy followed parallel transects perpendicular to the coast with continuous acoustic data recorded to a maximum depth of 500 meters, being the data processed by echointegration per cells of 0.1 nm long and 5 m height (**Figure 1**; **Figure S1**). Fishing hauls were conducted to validate species composition and to obtain length distribution. The acoustic echointegrations were allocated to species according to catch composition of the trawls and summed over the entire water column, resulting in acoustic abundance values referred to as NASC (Nautical Area Scattering Coefficient; m2nm-2) (**Figure 2a**).

**Figure 1.**
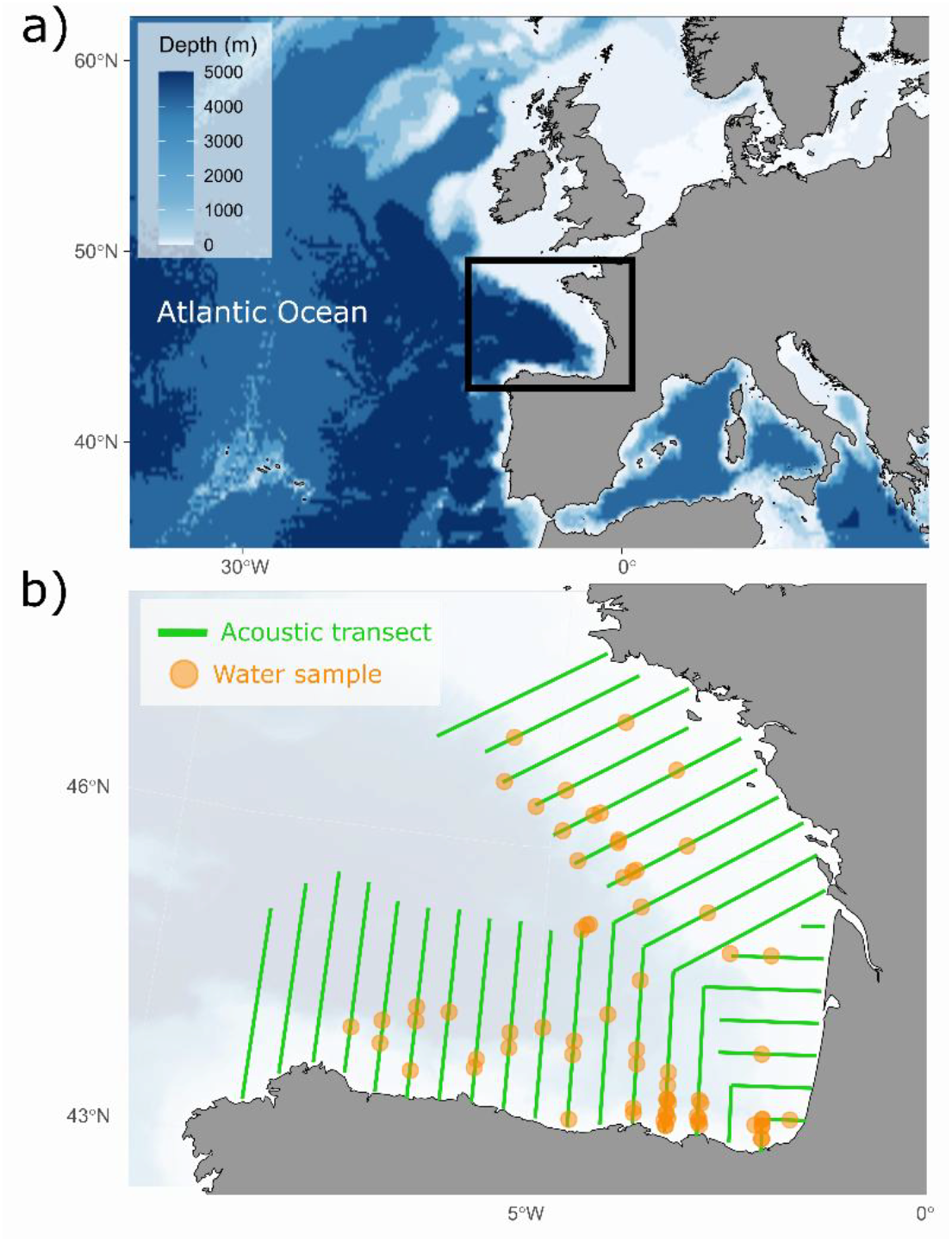
Area of study. **a)** Location of the Bay of Biscay in the Northeast Atlantic Ocean. **b)** Acoustic transects (green) and eDNA sampling stations (orange) covered in the JUVENA scientific surveys during the 6-year period covered in this work.

**Figure 2.**
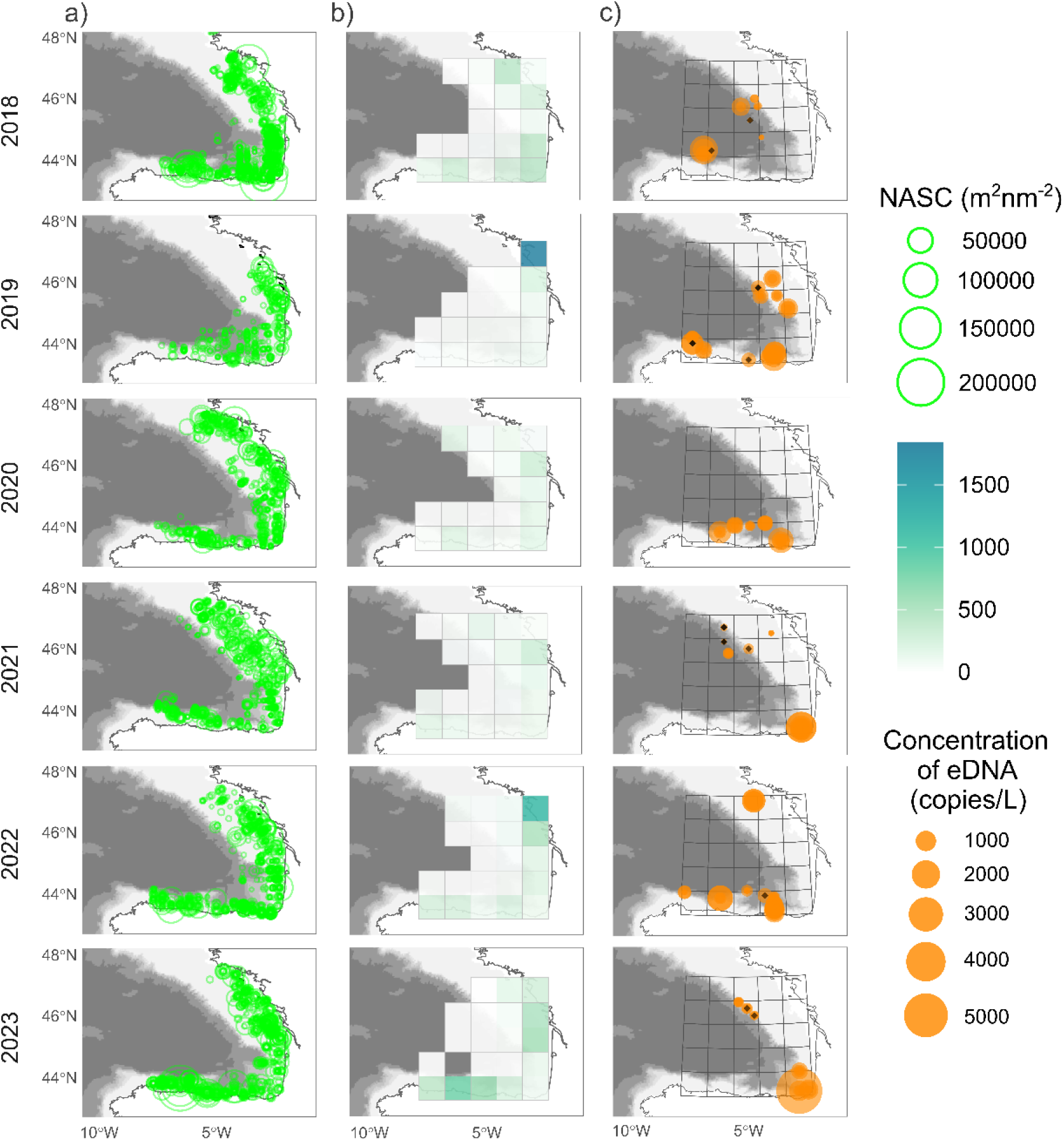
Anchovy detections in the BoB during the 6-year-period. **a)** Depth-integrated acoustic-trawl derived signal (NASC) assigned to anchovy per cell. **b)** Acoustic signal grided by area. c) Distribution and abundance of anchovy eDNA per sample and grided area showing to which grid each group of samples corresponds. Black diamonds represent samples where no anchovy eDNA was detected.

### eDNA sample collection and extraction

In parallel to acoustic sampling, water samples were collected during the 2018-2023 JUVENA surveys, resulting in a total of 255 seawater samples distributed among 70 stations (**Figure 1**; **Figure S1**). 4-5 litre water samples were collected at each station using a CTD rosette with Niskin bottles and filtered through 0.45 μM Sterivex enclosed filters (Milipore). Filters were stored onboard at -20ºC. DNA was extracted from the filters using DNeasy Blood & Tissue Kit (Qiagen, USA), following the modifications described in Spens *et al*. (2017). The concentration of the total extracted DNA (μg/ml) was calculated with fluorimetry (Qubit™), its quality through UV spectrometry (NanoDrop™) and its integrity through agarose gel electrophoresis. Original eDNA samples were diluted to a concentration of 5 ng µl −1 for subsequent analyses.

### Digital PCR (dPCR) analysis

Water samples were amplified using a QIAcuity Digital PCR system (Qiagen), with a species-specific quantitative method to estimate absolute concentrations of target DNA in seawater. For the detection of European anchovy we used the assays developed by Albaina *et al*. (2015). Because the assay was originally designed for qPCR, we optimized the assay to be run in digital PCR (dPCR) and the species-specific fluorescent TaqMan probe was labelled with FAM. Each reaction consisted of 2 µl of template DNA, 10 µl PCR Master Mix, 3.2 µl of each target-specific probe and primers (5 uM) and 18.4 µl water, resulting in a total reaction volume of 40 µl. Reaction mixes were transferred into the QIAcuity 24-well Nanoplates (maximum 26,000 partitions per reaction). Each plate included a positive control (DNA extracted from anchovy tissue sample) and one negative control (no template). The assay was amplified in the QIAcuity following these conditions: an initial step at 95°C for 2 minutes, followed by 40 cycles alternating between 95°C for 15 seconds and 60°C for 60 seconds. Fluorescence was measured and the number of eDNA copes per reaction volume were calculated by the QIAcuity Software Suite software (version 2.1.7.). The results were visualized using scatterplots created by the software and the positive threshold was manually determined based on the highest droplets in the negative controls of each plate. The software calculates the average number of molecules per partition using Poisson statistics and then transforms this information into concentration (Cr) in the reaction by dividing by partition volume as follows:

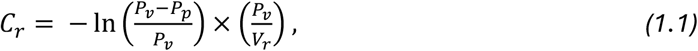

where *Pv* corresponds to valid partitions, *Pp* refers to positive partitions and Vr is the reaction volume (i.e., 40 µl). The total effective concentration (CL) (i.e., target eDNA copies per litre of sea water) was calculated for each sample using the formula:

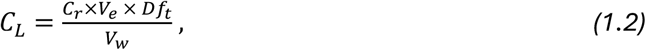

where Cr is copies of target eDNA per microlitre, Ve is the total elution volume after extraction (in ml), Vw is the volume of filtered sea water (in litres) and Dft is the unitless dilution factor, calculated as:

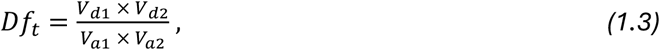

where Vd1 is the volume of the first dilution (i.e., the aliquot), Va1 is the volume of the elution used to create the aliquot, Vd2 is the volume of the second dilution (dPCR reaction) and Va2 is the volume of the aliquot used in the dPCR reaction. The final concentration values were calculated as copies of target DNA per litres of seawater.

### Spatial gridding of both datasets and acoustic-trawl data resizing

To obtain combinable datasets, we gridded the sampling area horizontally in ∼100×150 nm length resolution cells (i.e., 0.8×1.2 degrees of latitude and longitude, respectively) and we determined the trawl-acoustic and eDNA data that corresponded to each spatial unit by identifying the samples found in each cell according to their geographic coordinates. Trawl-acoustic data were segmented according to cells and the mean of all depth-integrated values located in each cell was calculated per year. Acoustic sampling effort covered the study area completely, resulting in 120 grided cell values (**Figure S2a**). The eDNA samples were not spatially merged and this information was used as point data. A total of 29 cells had eDNA data, with from 3 to 12 point values (average = 10) over the 6-year period (**Figure S2b**).

### Description and application of the spatial joint model

We developed a Bayesian model to integrate both acoustic-trawl and eDNA data to jointly estimate the underlying fish biomass as a latent parameter. We assumed that both acoustic and eDNA data provide imperfect independent information about the underlying true abundance of fish, and that each data type observation is linearly and positively correlated with fish biomass (i.e., the bigger the true fish abundance, the stronger the acoustic signal and the greater the concentration of eDNA).

### Observation models

We modelled the logarithm of the observed acoustic signal intensity (i.e., NASC), Wi, of anchovies at gridded area *i* as drawn from a normal distribution with a mean species density, θi, and a standard deviation φ:

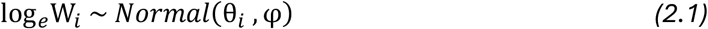

Likewise, we modelled the logarithm of the observed eDNA density (i.e., copies DNA/L seawater), Yi, with a normal distribution with a mean concentration, µi, and a standard deviation σ*i*:

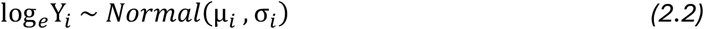

Note that the standard deviation for eDNA was allowed to vary by gridded area because point data is used, while for acoustics it was assumed to be constant in space due to integrated values. A constant value (1) was added to all eDNA data before transforming to the logarithmic scale to obtain valid values for the null observations (i.e., 0 copies/L). Also, all years of data were simultaneously treated because it was assumed that the spatial distribution of anchovies did not change over time, considering that sampling took place consistently during late early autumn.

### Process models

The observations from both methods were then related to a shared reality, that is, the underlying fish abundance. Let Xi be the true – but unobserved – biomass (in log scale) of anchovies in grid cell *i*. The expected means of the lognormal distributions of the observations were linked linearly to the underlying fish biomass (Xi).

Specifically, the mean of the acoustic observation, θi, of anchovies at grid cell *i* was modelled as a linear function,

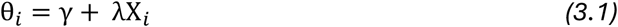

and the mean of the eDNA abundance, µi, as a linear process,

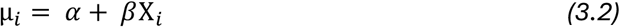

where α and γ are the intercepts, and β and λ are the coefficients of Xi. Likewise, the standard deviation of the eDNA estimations (σi, which is constrained to be a positive value) was an exponential linear function of the fish biomass (Xi),

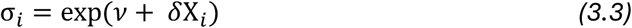

where *ν* is the intercept and δ the coefficient of Xi.

The model was fitted in a Bayesian framework using the prior distributions detailed in **Table S1**. The sensitivity of the model results to the priors was evaluated by running the model with different prior distributions. The joint model results remained consistent across the different prior distribution runs, indicating that they were not strongly affected by prior parameter values (**Figure S3**) and that the model’s outputs are robust, being driven by the data itself, rather than by initial assumptions.

All the analyses were conducted in Rstudio (version 4.2.2) as an integrated development environment for R (R Core Team 2022. The model was coded and fitted using the package *rstan* (Bürkner 2017) that is based on the Stan programming language. Stan is written in C++ and implements full Bayesian statistical inference using Hamiltonian Markov Chain Monte Carlo (MCMC) (Carpenter *et al*. 2017). The joint model was run with a step size of 0.5 and 4 chains, each with 4000 sampling iterations, including a burn-in of half of the total iterations and thinning of every sample. The model convergence was checked through the R-hat convergence diagnostic (Vehtari *et al*. 2021) and by visual examination of the resulting correlation plots and chain mixtures. The posterior distribution of the parameters showed that the Markov chains effectively converged (R-hat ≈ 1), indicating that the model sampling process was stable and consistent. The effective number of independent samples ranged between 1000-12000, which ensures reliable estimates (**Figures S4**).

Finally, the posterior distributions of the biomass in logarithmic scale (Xi) were used to estimate spatial abundance patterns for anchovy in the BoB. The Stan model developed scripts and corresponding output files are available at GitHub (https://github.com/cclaver001/IMPACT).

### Evaluation of the contribution of eDNA to the joint model

One of the aims of this work was to determine the contribution of eDNA-derived data to the improvement of the estimates when used together with acoustic data. In order to evaluate the marginal value of eDNA, we extracted and ran only the trawl-acoustic section of the model with the same procedure (hereby referred to as acoustic model) (**Table S2**) and compared the change in the estimates between the joint model and the acoustic model by calculating the difference of the biomass in logarithmic scale (Xjoint - Xacoustic). The acoustic model converged as well as the joint model (i.e., R-hat values close to 1, large effective sample size and well-mixed chains) (**Figure S5**).

## Results

### Acoustic and eDNA based anchovy abundance data

A maximum of 22 and a minimum 18 grided areas had acoustic signal for each of the 6 years, with a total of 120 grided areas considered. From them, 80% were positive for anchovy. As expected in light of the known anchovy distribution (Boyra *et al*. 2013), the acoustic signal (average NASC = 157 m2nm-2) was higher in the cells adjacent to coast than in those representing more oceanic areas, with the highest acoustic signal (≈1800 m2nm-2) detected in the northern coastal French shelf (**Figure 2b**).

Regarding the eDNA samples, 93% of the eDNA samples resulted positive for anchovy, with an average value of 470 copies/L and the highest value being 5800 copies/L (**Figure 2c**). Most of the samples were collected between the surface and 300 m depth, being the highest DNA concentrations of anchovy DNA found in the first hundred meters of the water column with a decreasing concentration with depth (**Figure S6**). In some stations, anchovy eDNA was detected in deeper areas than normally expected, which might be related to sinking, given the positive detection in the same area in shallower waters (Harrison *et al*. 2019). Also, higher DNA concentrations were found in close-to-shore and shallow waters than in oceanic waters, far from shore (**Figure S8**), matching the known distributions and preferences of anchovies in the area (Boyra *et al*. 2013).

### Performance of the joint model

Parameters referring to intercepts (alpha, gamma) and slopes (beta, lambda) of the relationship between eDNA and acoustic data with biomass had narrow credible intervals that indicated high precision (**Table S4**). Fish biomass estimates (*X*) had the widest intervals (i.e., greatest variability), likely due to differences within sites, sampling effort or inherent biological variability. Grid cells with higher biomass estimates had systematically lower standard deviations, indicating greater precision. Analysis of the posterior parameter correlations provided further insight into the model’s behaviour and the interdependencies among parameters (**Figure S5**). A positive correlation between alpha and gamma indicates that sites with high eDNA mean values also tend to have high acoustic mean values. This correlation suggests that the model is capturing meaningful relationships between acoustic and eDNA signal, as assumed, being both positively linked to biomass: the more biomass, the more acoustic and eDNA signal.

A negative correlation between alpha and nu indicates a trade-off in how the eDNA data fits the model: a higher mean eDNA is compensated by a lower estimated variability, and vice versa, to best explain the observed dispersion of eDNA data points, particularly where fish biomass is low. Similarly, the negative correlation between gamma and nu suggests an interdependency between the estimated acoustic signal and the estimated variability in eDNA measurements. This arises because the model jointly estimates parameters for both data sources and the biomass; a stronger acoustic signal might help constrain overall uncertainty, allowing the model to estimate lower variability in the eDNA estimates. Crucially, these correlations align well with the common understanding that higher biomass values might be measured with greater precision, whereas lower values might be more susceptible to variability due to natural stochasticity or noise (Hespanhol *et al*. 2019). Such posterior parameter interdependencies are a common feature of complex statistical models, especially those involving multiple observation processes linked by shared latent variables, such as the one described in this work.

### Biomass estimates of the joint model and differences from the acoustic model

Overall, the joint model linked the information derived from two independent observation methods and provided biomass estimates of anchovies, in units of fish per grided area (**Figure 3a**). In contrast, the acoustic model used only acoustic data to estimate anchovy biomass (**Figure S9**), and the results indicated, consistent with the acoustic signal, that anchovy biomass is mostly located in the continental shelf, concentrated in the areas closest to the shore.

**Figure 3.**
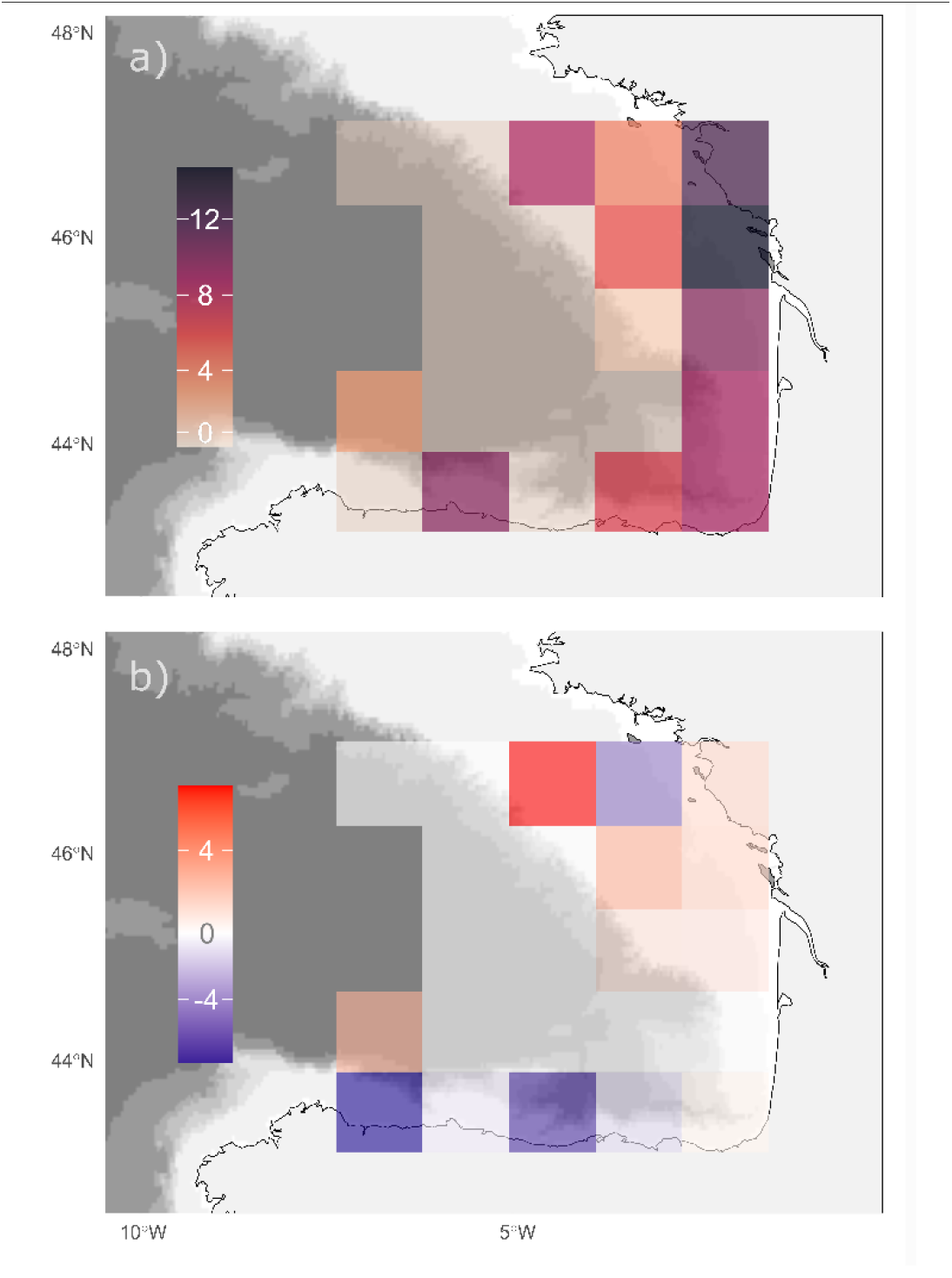
Distribution and abundance of European anchovy in the BoB for the 6-year period. **a)** Posterior estimates of anchovy biomass estimates (mean) under the joint model, representing depth-integrated biomass in logarithmic scale. Negative values in logarithmic scale (corresponding to low biomass levels) were replaced by zero. **b)** Change in posterior estimates of the depth-integrated biomass (mean values of biomass) when applying the joint model, compared to the only acoustic version of the model (X_joint_ - X_acoustic_).

Largely, both acoustic and eDNA data showed similar spatial patterns and, consequently, the two models agreed in about half of the gridded areas (i.e., zero and low values in **Figure 3b**). In some gridded areas, instead, eDNA and acoustic data disagreed, and the estimates differed among the models. For instance, the joint model estimated lower anchovy abundance than the acoustic model in two cells along the Cantabrian Sea (i.e., negative values in **Figure 3b**). This discrepancy stems from the eDNA data, where positive anchovy values were detected in those cells for only one year over the 6-year period, and the joint model diluted the perceived presence of anchovy in those areas. Conversely, according to the joint model, anchovies showed a wider spatial distribution along the continental shelf in both the Spanish and French waters, with a progressive decay in abundance in deeper waters and offshore (**Figure 3a**). Compared to the acoustic model, this results in higher values for anchovy in these areas (i.e., positive values in **Figure 3b**).

Thus, compared to the acoustic model, the primary contribution of this approach lies in the model’s capacity to synthesize information from two independent observation methods: linking acoustic and eDNA data to a shared latent variable representing true biomass, the joint model leverages their combined use, offering a more comprehensive view than either method alone.

## Discussion

### Towards an effective integration of eDNA for providing assessment-informative data

Despite the high anticipation surrounding the use of eDNA in fisheries (Gilbey *et al*. 2021), there have been limited efforts to propose methods for integrating eDNA data into assessment and translating it into practical information. The independent use of eDNA data to develop a biomass index, as evidenced by Shelton *et al*. (2022), represents a significant accomplishment that underscores the value and potential of this information source.

Although oceanographic surveys are currently the best opportunity to collect eDNA broadly in marine areas, eDNA sampling efforts are generally limited during campaigns because they are mostly opportunistic, and datasets of fine scale covered study areas like the one obtained by Shelton *et al*. (2022) are difficult to obtain. In our study, the opportunistic annual eDNA sampling was limited and did not cover the sampling area evenly. To address this, we merged the time series data, but some gaps remained in some areas. Alternatively, Bayesian joint models are a reliable approach for combining eDNA information with other fisheries data, such as trawls, as demonstrated by Guri *et al*. (2024). Therefore, we decided to combine eDNA with acoustic-trawl data in a Bayesian joint model, resulting in the first integration these two data sources to produce anchovy biomass estimates in space.

Different data sources offer varying pieces of information which, when combined, can provide more comprehensive and accurate insights. For example, catches and acoustic data represent snapshots of a certain moment in a given space, resulting in discrete and heterogeneous records. Instead, eDNA is composed of DNA molecules from fish-derived particles, that create a more uniformly distributed scattered signal in the environment. Accordingly, our results showed that high concentrations of anchovy eDNA were more widespread than acoustic signal detections and, accordingly, the joint model estimated higher anchovy abundances in a wider distribution area than the acoustic-only model estimates. This can be explained due to vessel avoidance in challenging sampling areas, such as shallow coastal waters (Draštík and Kubečka 2005, De Robertis and Handegard 2013), which can lead to acoustic derived biomass underestimations. Additionally, when adult anchovies are near rocky bottoms, trawling is not feasible and delayed night trawls are used (Boyra et al. 2016), making it harder to identify acoustic signals. Our results suggest that eDNA analysis could potentially address false absences of anchovy derived from the acoustic data, thereby making biomass derived from joint estimates more representative and accurate.

### Challenges and strategies for the use of eDNA in stock assessment

Identifying key milestones for integrating eDNA data into fisheries assessment will help focus future efforts. Although improvements like data modelling could facilitate eDNA information integration into stock assessments, some constraints must be considered.

For example, the data for stock assessment derived from the JUVENA survey only include calculations based on juveniles, and current eDNA analyses fail at differentiating among life stages. Thus, understanding the species’ biology is critical to properly interpret at eDNA data; for instance, with adults remaining close to the bottom and juveniles forming pelagic shoals near the surface during daytime (Boyra *et al*. 2016), it is reasonable to assume that the eDNA collected is predominantly from juvenile individuals, as the sampling was conducted during daylight hours in late early autumn. In the future, using epigenetics or environmental RNA, which can provide information about a specific group of individuals of a species, could also help to solve this issue (Stevens and Parsley 2023).

An additional challenge for eDNA-based studies is that eDNA sampling is often opportunistic and sampling effort is significantly smaller than for other methods (e.g., 37 water samples vs 141 trawls (Le Joncour *et al*. 2024)). This may lead to significant variations in dataset sizes, leading to estimates biased towards the most available data when models are applied. This is likely to happen in scientific surveys that are focused on a specific methodology, such as JUVENA, where the acoustic effort is considerably bigger than the eDNA effort. Indeed, incorporating a new sampling methodology into a well-established survey can be challenging, especially when the nature of the new method is different. For instance, in our case, the acoustic data is collected continuously across transects whereas eDNA is sampled in point-stations, so that the vessel must interrupt the acoustic sampling to deploy the Niskin bottles. In recent years, automatic eDNA samplers have been developed (George *et al*. 2024) and can represent a promising sampling strategy to combine eDNA and acoustic sampling onboard. However, passive eDNA sampling would only be feasible to collect water from the surface through continuous water intake systems of the vessels, which might not be as representative as vertical profile sampling. Given that most pelagic species make nocturnal vertical migrations to the surface, this could be overcome by carrying out the continuous sampling during the night, as in (Shelton *et al*. 2022).

On the other hand, one of the major limitations of this work is the integration of data from a time-series, as annual estimates are needed for evaluation purposes if the results are to be used directly. We have combined data from several years due to the limited per-year eDNA sampling coverage. However, this is a pilot study that is expected to lay the foundation for a consistent sampling in the future that will allow repeating these analyses for each year to come. Ideally, the eDNA effort should be increased (Cote *et al*. 2023) and a proper sampling design that could be repeated and be consistent along time should be established. One way for avoiding weakening the eDNA information due to the lower sample size is to weight data differently when using the models. In this work, we have reduced the acoustic data to cells and maximised the eDNA information by using all water samples separately, without combining them. Because the eDNA sampling locations and effort was variable across the 6-year period, the size of the grids used in this study was determined to obtain at least one eDNA value per most of the gridded areas, ensuring a consistent coverage across the study region. However, a bigger sampling effort with a larger spatial coverage would allow to draw smaller cells and could be an interesting advancement in future surveys to obtain more detailed maps. In sum, identifying these improvements provides a roadmap for future developments based on work such as that described in this manuscript.

## Conclusion

Here, we have successfully combined anchovy acoustic-trawl and eDNA based abundance data and show that the resulted joint distribution and abundance patterns obtained are consistent with the current knowledge of anchovy in the BoB. The versatility of Bayesian models allows the joint model proposed here to be used with different input data (other than eDNA and acoustic trawl) and species, which increases its potential use in other case studies. Also, this work exemplifies how eDNA data can be used as quality control of acoustic indices, detecting potential catchability problems, and thus helping to improve the robustness and precision of the acoustic estimates. This integrated approach not only utilizes the complementary nature of the data but also provides valuable insights into the relationship between acoustic signals, eDNA concentrations, and biomass.

## Acknowledgements

Authors are grateful to the crew of RR/VV Angeles Alvariño and Emma Bardán, specially to Carlota Pérez, Naiara Serrano and Deniz Kukul, for their support during sample collection, and to Aitor Lekanda and Iosu Paradinas, for helping with the acoustic data interpretation and for providing feedback on the model, respectively. The authors would like to thank “The eDNA Collaborative” team, and particularly Andrew Olaf Shelton, for providing the foundations on which current work lies, as well as relevant input throughout the development of the project. The JUVENA survey is co-funded by the Department of Economic Development, Sustainability and the Environment, by the Vice-Ministry of Agriculture, Fisheries and Food Policy of the Directorate of Fisheries and Aquaculture of the Basque Government, as well as by the Spanish Institute of Oceanography of the Ministry of Science and Innovation and by the General Secretariat of Fisheries of the Ministry of Agriculture, Fisheries and Food (MAPA) of the Spanish Government. This work has been funded by the European Union’s Horizon 2020 research and innovation program (project OBAMA-NEXT with grant agreement No. 101081642, project BIOcean5D with grant agreement No. 101059915) and the Department of Education (Basque Government) through a predoctoral grant to Cristina Claver.

## Author contribution

Conceptualization: C.C., R.P.K. Formal analysis: C.C., R.P.K. Funding acquisition: C.C., G.B., N.R-E. Investigation: C.C., B.S., O.C., I.M., G.B., R.P.K., Methodology: C.C., R.P.K., Resources: B.S., G.B., N.R-E. Software: C.C., L.I., R.P.K. Validation: L.I. Visualization: C.C. Writing – original draft: C.C., R.P.K, N.R-E. Writing – review & editing: all authors.

## Data Availability Statement

The data underlying this article are available in GitHub and can be accessed at https://github.com/cclaver001/IMPACT.

## References

Albaina A, Irigoien X, Aldalur U, et al. A real-time PCR assay to estimate invertebrate and fish predation on anchovy eggs in the Bay of Biscay. Progress in Oceanography 2015; 131: 82–99.

Barange M, Bernal M, Cergole MC, et al. Current trends in the assessment and management of stocks. In: Checkley D, Alheit J, Oozeki Y, et al. (eds.), Climate Change and Small Pelagic Fish, Cambridge: Cambridge University Press, 2009, 191–255

Borja A, Amouroux D, Anschutz P, et al. The bay of biscay. In: World Seas: An Environmental Evaluation, Elsevier, 2019, 113–152

Boyra G, Cotano U, Martinez U, et al. JUVENA series review of the spatial distribution of anchovy juveniles in the Bay of Biscay. In: XI International Symposium on Oceanography of the Bay of Biscay, San Sebastián (Spain), 2008

Boyra G, Martínez U, Cotano U, et al. Acoustic surveys for juvenile anchovy in the Bay of Biscay: abundance estimate as an indicator of the next year’s recruitment and spatial distribution patterns. ICES Journal of Marine Science 2013; 70: 1354–1368.

Boyra G, Peña M, Cotano U, et al. Spatial dynamics of juvenile anchovy in the Bay of Biscay. Fisheries Oceanography 2016; 25: 529–543.

Boyra G, Sobradillo B, Rico I, et al. Acoustic surveying of anchovy juveniles in the Bay of Biscay: Juvena 2023 survey report. In: Technical Report, Working Document to ICES WGACEGG meeting, Lisboa, Portugal …, 2023

Bürkner P-C. brms: An R package for Bayesian multilevel models using Stan. Journal of statistical software 2017; 80: 1–28.

Carpenter B, Gelman A, Hoffman MD, et al. Stan: A probabilistic programming language. Journal of statistical software 2017; 76.

Checkley Jr DM, Asch RG, Rykaczewski RR. Climate, anchovy, and sardine. Annual Review of Marine Science 2017; 9: 469–493.

Chevrinais M, Bourret A, Côté G, et al. Improving an endangered marine species distribution using reliable and localized environmental DNA detections combined with trawl captures. Scientific Reports 2025; 15: 1–13.

Cote D, McClenaghan B, Desforges J, et al. Comparing eDNA metabarcoding and conventional pelagic netting to inform biodiversity monitoring in deep ocean environments. ICES Journal of Marine Science 2023; 80: 2545–2562.

De Robertis A, Handegard NO. Fish avoidance of research vessels and the efficacy of noise-reduced vessels: a review. ICES Journal of Marine Science 2013; 70: 34–45.

Doray M, Boyra G, van der Kooij J. ICES survey protocols–Manual for acoustic surveys coordinated under ICES working group on acoustic and egg surveys for small pelagic fish (WGACEGG). ICES techniques in marine environmental sciences 2021; 64.

Draštík V, Kubečka J. Fish avoidance of acoustic survey boat in shallow waters. Fisheries Research 2005; 72: 219–228.

Eichmiller JJ, Miller LM, Sorensen PW. Optimizing techniques to capture and extract environmental DNA for detection and quantification of fish. Molecular ecology resources 2016; 16: 56–68.

FAO FAOotUN. The State of World Fisheries and Aquaculture. In: action. BTi (ed.), Rome, 2024

Fleischer G. The 2003 integrated acoustic and trawl survey of Pacific hake, Merluccius productus, in US and Canadian waters off the Pacific coast. 2005.

Fraija-Fernández N, Bouquieaux MC, Rey A, et al. Marine water environmental DNA metabarcoding provides a comprehensive fish diversity assessment and reveals spatial patterns in a large oceanic area. Ecology and Evolution 2020; 10: 7560–7584. 10.1002/ece3.6482

Fréon P, Cury P, Shannon L, et al. Sustainable exploitation of small pelagic fish stocks challenged by environmental and ecosystem changes: a review. Bulletin of marine science 2005; 76: 385–462.

George SD, Sepulveda AJ, Hutchins PR, et al. Field Trials of an Autonomous eDNA Sampler in Lotic Waters. Environmental Science & Technology 2024.

Gilbey J, Carvalho G, Castilho R, et al. Life in a drop: Sampling environmental DNA for marine fishery management and ecosystem monitoring. Marine Policy 2021; 124: 104331. 10.1016/j.marpol.2020.104331

Guri G, Shelton AO, Kelly RP, et al. Predicting trawl catches using environmental DNA. ICES Journal of Marine Science 2024: fsae097.

Harrison JB, Sunday JM, Rogers SM. Predicting the fate of eDNA in the environment and implications for studying biodiversity. Proceedings of the Royal Society B 2019; 286: 20191409.

Hespanhol L, Vallio CS, Costa LM, et al. Understanding and interpreting confidence and credible intervals around effect estimates. Brazilian journal of physical therapy 2019; 23: 290–301.

ICES. Bay of Biscay and the Iberian Coast ecoregion – Fisheries Overview. In, 2022

Karlsson E, Ogonowski M, Sundblad G, et al. Strong positive relationships between eDNA concentrations and biomass in juvenile and adult pike (Esox lucius) under controlled conditions: Implications for monitoring. Environmental DNA 2022; 4: 881–893.

Klymus KE, Richter CA, Chapman DC, et al. Quantification of eDNA shedding rates from invasive bighead carp Hypophthalmichthys nobilis and silver carp Hypophthalmichthys molitrix. Biological conservation 2015; 183: 77–84.

Knudsen SW, Ebert RB, Hesselsøe M, et al. Species-specific detection and quantification of environmental DNA from marine fishes in the Baltic Sea. Journal of experimental marine biology and ecology 2019; 510: 31–45.

Lacoursière-Roussel A, Côté G, Leclerc V, et al. Quantifying relative fish abundance with eDNA: a promising tool for fisheries management. Journal of Applied Ecology 2016; 53: 1148–1157.

Le Joncour A, Mouchet M, Boussarie G, et al. Is it worthy to use environmental DNA instead of scientific trawling or video survey to monitor taxa in soft-bottom habitats? Marine Environmental Research 2024; 200: 106667.

Maes SM, Desmet S, Brys R, et al. Detection and quantification of two commercial flatfishes (Solea solea and Pleuronectes platessa) in the North Sea using environmental DNA. Environmental DNA 2024; 6: e426.

Maiello G, Talarico L, Carpentieri P, et al. Little samplers, big fleet: eDNA metabarcoding from commercial trawlers enhances ocean monitoring. Fisheries Research 2022; 249: 106259.

Salter I, Joensen M, Kristiansen R, et al. Environmental DNA concentrations are correlated with regional biomass of Atlantic cod in oceanic waters. Communications Biology 2019; 2: 461.

Sánchez S, Ibaibarriaga L, Uriarte A, et al. Challenges of management strategy evaluation for small pelagic fish: the Bay of Biscay anchovy case study. Marine Ecology Progress Series 2019; 617: 245–263.

Shelton AO, Ramón-Laca A, Wells A, et al. Environmental DNA provides quantitative estimates of Pacific hake abundance and distribution in the open ocean. Proceedings of the Royal Society B 2022; 289: 20212613.

Simmonds J, MacLennan DN. Fisheries acoustics: theory and practice. John Wiley & Sons, 2008

Spens J, Evans AR, Halfmaerten D, et al. Comparison of capture and storage methods for aqueous macrobial eDNA using an optimized extraction protocol: advantage of enclosed filter. Methods in Ecology and Evolution 2017; 8: 635–645.

Stenevik EK, Vølstad JH, Høines Å, et al. Precision in estimates of density and biomass of Norwegian spring-spawning herring based on acoustic surveys. Marine Biology Research 2015; 11: 449–461.

Stevens JD, Parsley MB. Environmental RNA applications and their associated gene targets for management and conservation. Environmental DNA 2023; 5: 227–239.

Stoeckle MY, Adolf J, Charlop-Powers Z, et al. Trawl and eDNA assessment of marine fish diversity, seasonality, and relative abundance in coastal New Jersey, USA. ICES Journal of Marine Science 2021; 78: 293–304.

Takahara T, Minamoto T, Yamanaka H, et al. Estimation of fish biomass using environmental DNA. PloS one 2012; 7: e35868.

Thomsen PF, Møller PR, Sigsgaard EE, et al. Environmental DNA from seawater samples correlate with trawl catches of subarctic, deepwater fishes. PloS one 2016; 11: e0165252.

Uriarte A, Ibaibarriaga L, Sánchez-Maroño S, et al. Lessons learnt on the management of short-lived fish from the Bay of Biscay anchovy case study: Satisfying fishery needs and sustainability under recruitment uncertainty. Marine Policy 2023; 150: 105512.

Vehtari A, Gelman A, Simpson D, et al. Rank-normalization, folding, and localization: An improved R for assessing convergence of MCMC (with discussion). Bayesian analysis 2021; 16: 667–718.

Veron P, Rozanski R, Marques V, et al. Environmental DNA complements scientific trawling in surveys of marine fish biodiversity. ICES Journal of Marine Science 2023; 80: 2150–2165.

Zhang J, Chen X, Zhou Q, et al. Species identification and biomass assessment of Gnathanodon speciosus based on environmental DNA technology. Ecological Indicators 2024; 160: 111821.

Zhou S, Fan C, Xia H, et al. Combined use of eDNA metabarcoding and bottom trawling for the assessment of fish biodiversity in the Zhoushan Sea. Frontiers in marine science 2022; 8: 809703.

