## supplementary material for "Integrating eDNA and acoustic-trawl data to provide small pelagic biomass estimates for fisheries assessment"

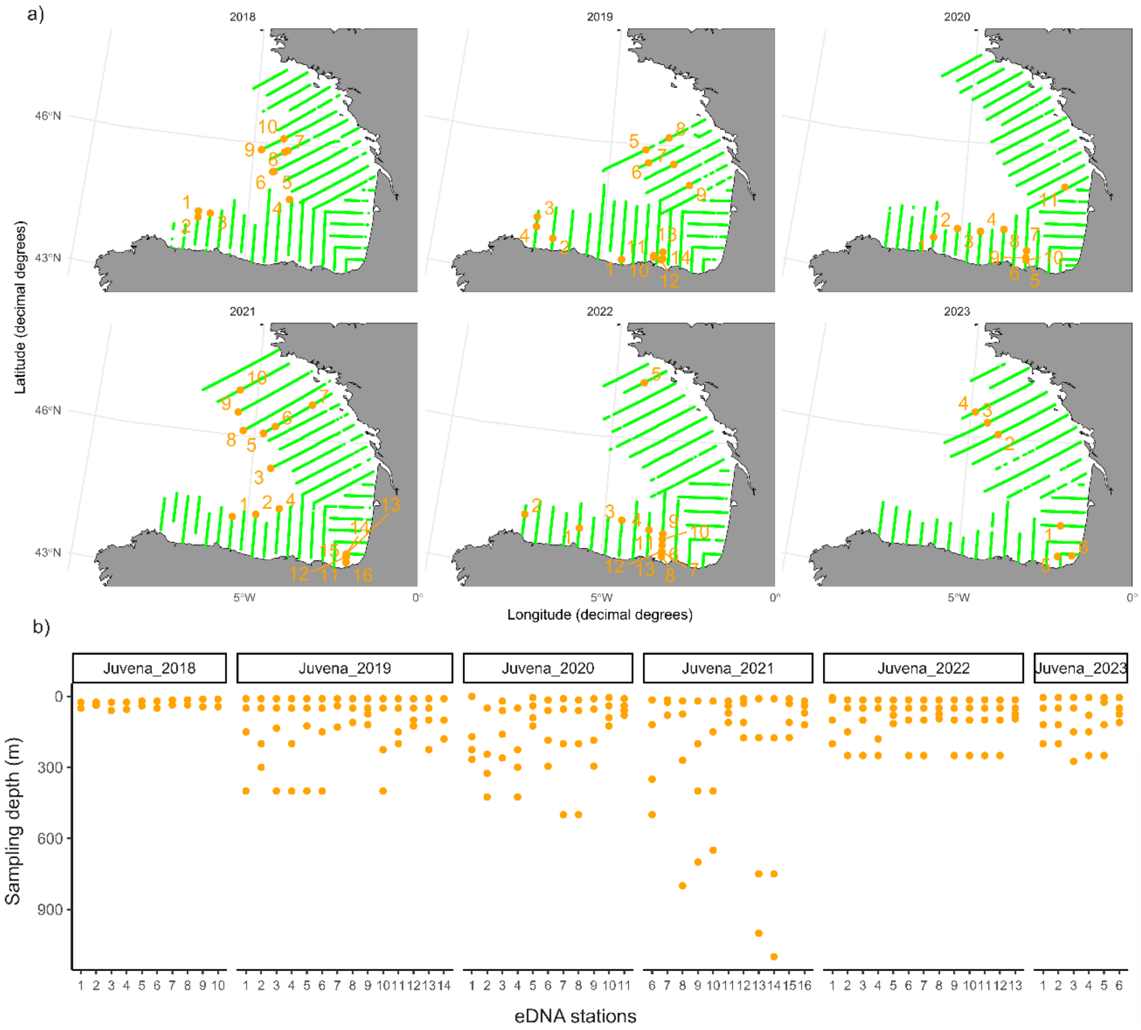


**Figure S1.** Sampling area. **a)** Facets represent the covered acoustic transects (green) and the eDNA stations (orange) for each year. **b)** Vertical sampling in the eDNA stations, each orange dots representing one sample.

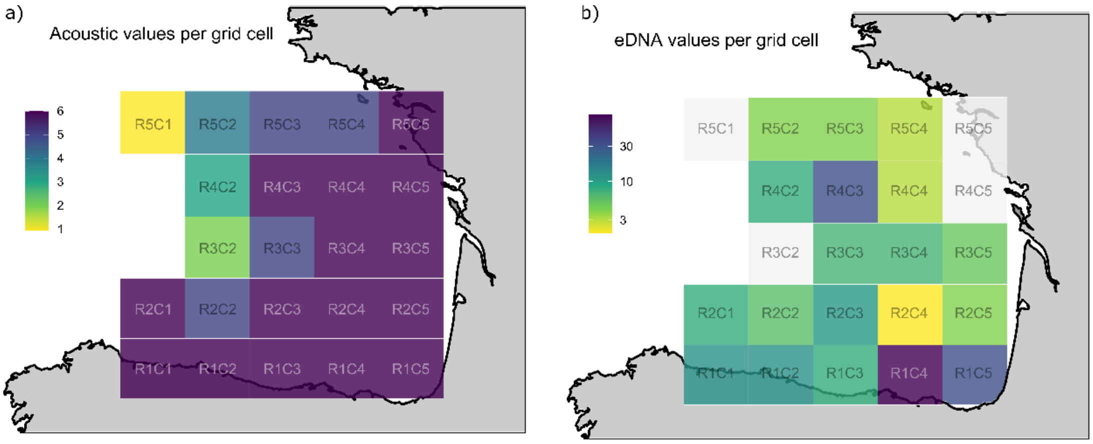


**Figure S2.** Grided map of the sampling area showing cumulative number of years with **a)** acoustic-trawl and **b)** genetic values for each cell. White cells have no genetic information. Names shown in the grid correspond to the identifiers of each cell to which the acoustic-trawl and genetic data are assigned.

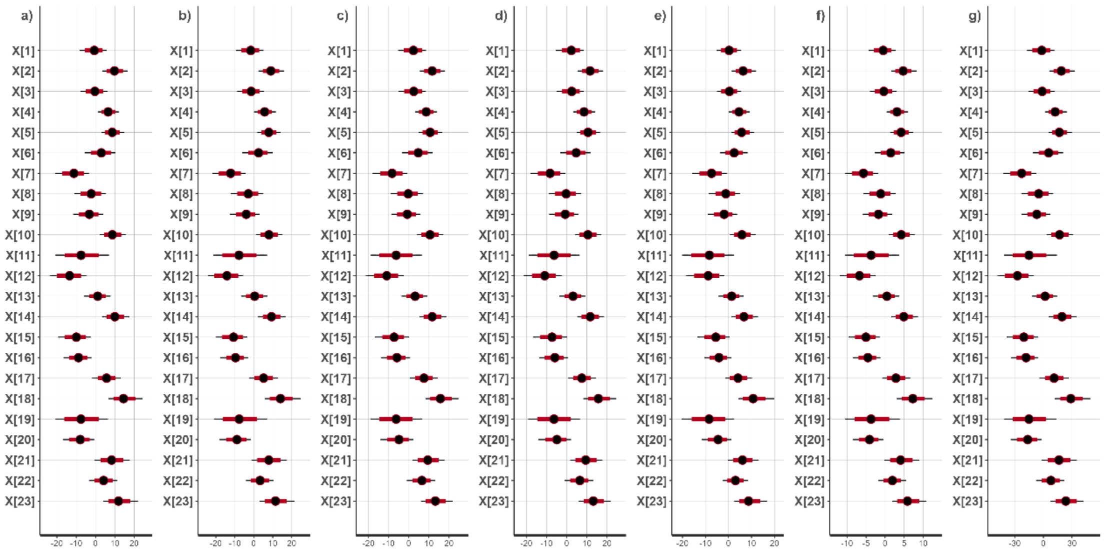


**Figure S3. Sensitivity test for parameter X with different priors. a)** Priors used in this study (see Table S2). **b)** Increased variance of normal priors (alpha ~ N(0,10); nu ~ N(-1,2); delta ~N(0,2); lambda ~N(0,10); gamma ~ N(-1,4)). **c)** Decreased variance of normal priors (alpha ~ N(0,2); nu ~ N(-1,0.5); delta ~N(0, 0.5); lambda ~N(0,2); gamma ~ N(-1,1)), **d)** Changed shape and rate parameters of gamma priors (beta ~G(2,2); sigma ~ (2,4); phi ~ G(0.5,0.5)), **e)** Use of extreme values (alpha ~N(0,50); beta ~ G(10,1); sigma ~ G(0.1, 1.0)), **f)** Reduced variance for X (X ~ N(0,5)) and **g)** increased variance for X (X ~ N(0,20)).

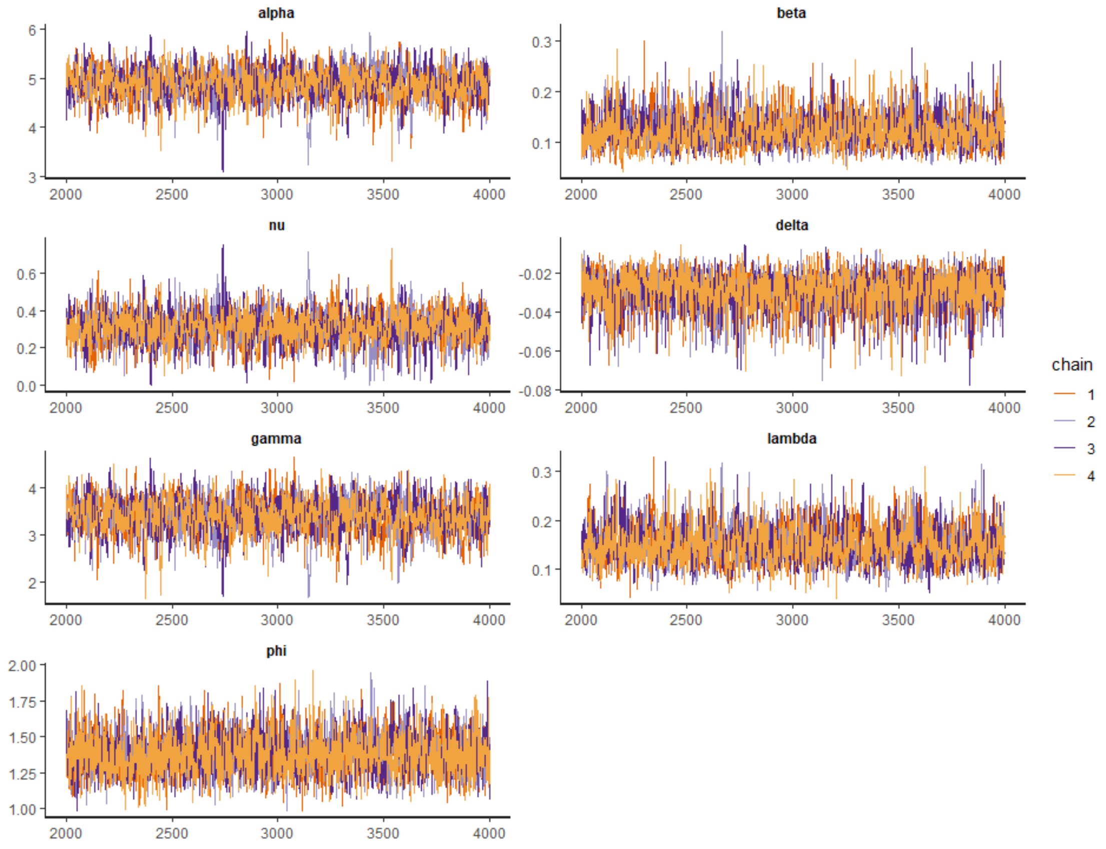


**Figure S4.** Markov chain trace plots showing the path for all chains of the parameters in the joint model.

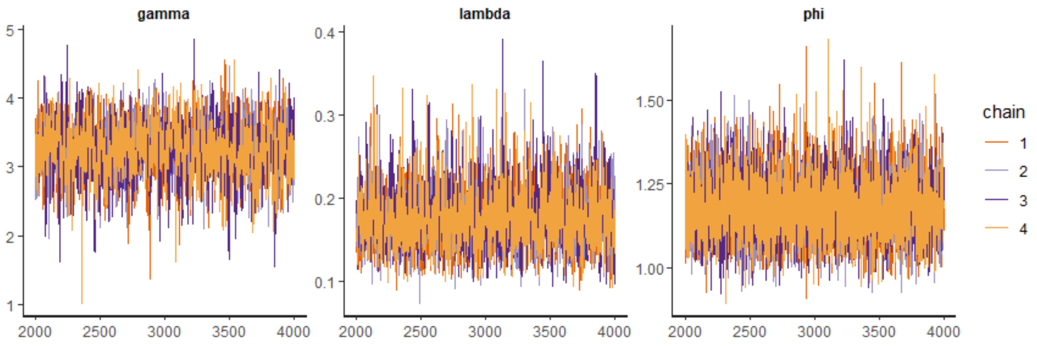


**Figure S5.** Markov chain trace plots showing the path for all chains of the parameters in the acoustic model.

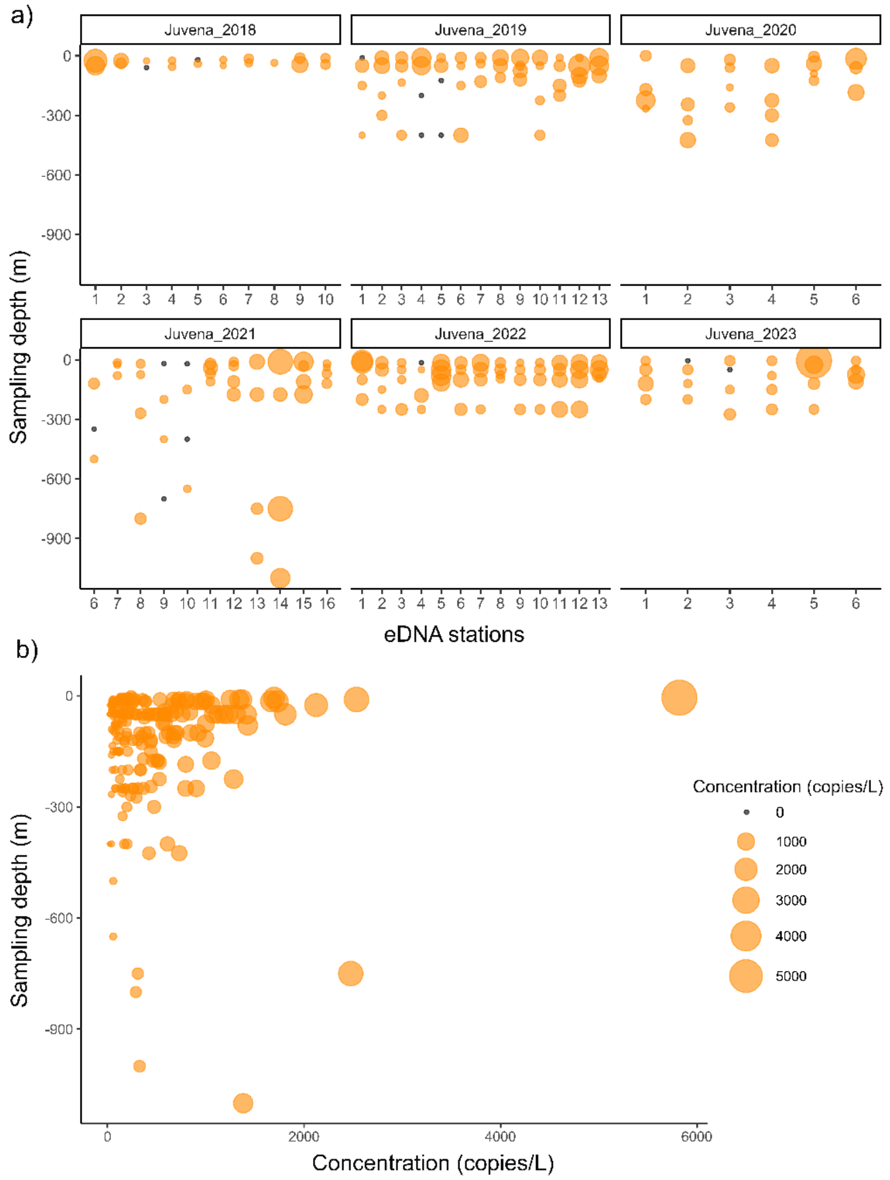


**Figure S6.** Concentration of DNA (copies/L) per point sample **a)** for each year and eDNA station and **b)** overall across the water column**.** Black points represent samples with no anchovy eDNA.

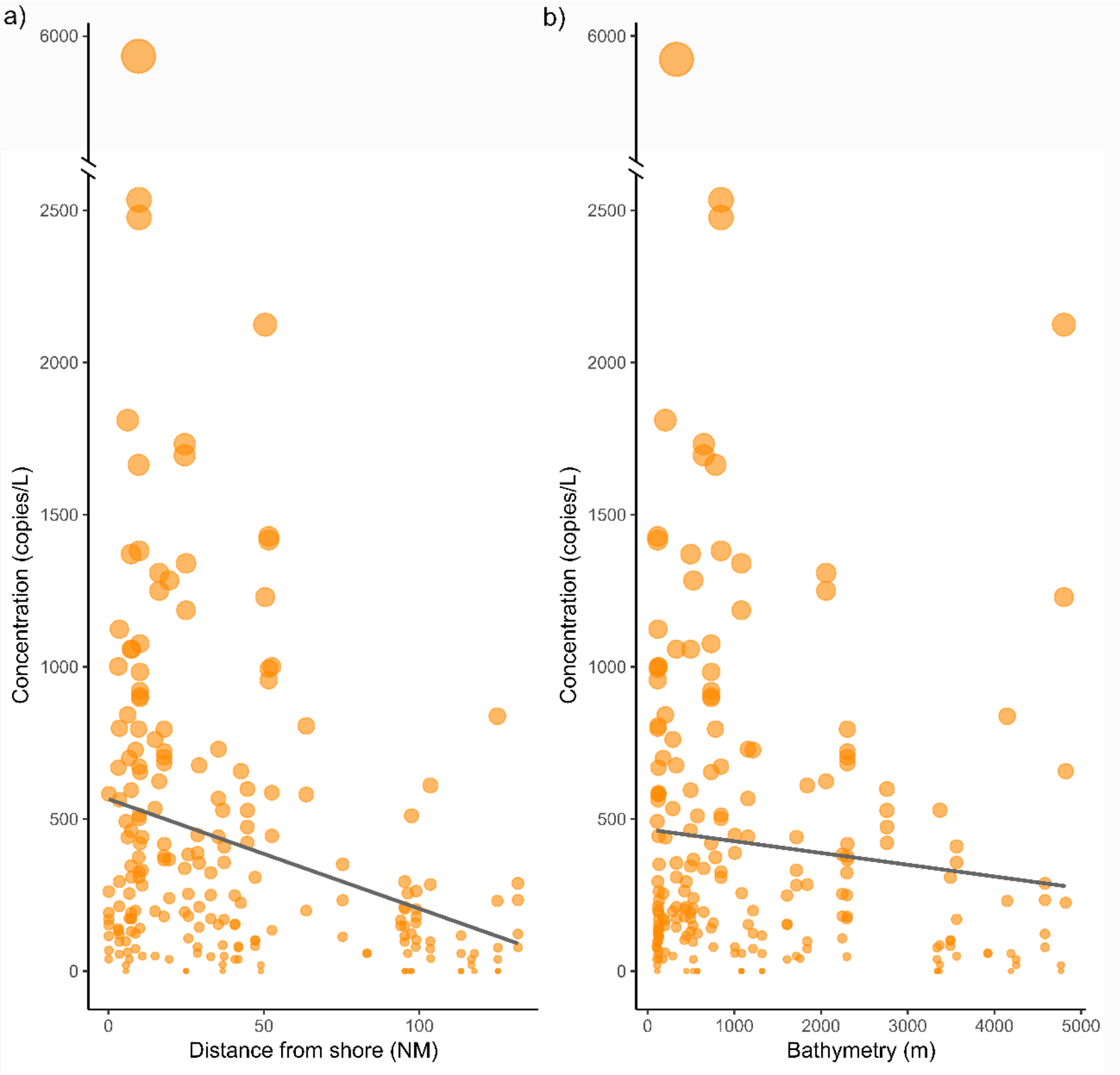


**Figure S7.** Linear relationship between concentration of DNA (copies/L) and **a)** distance from shore in nm or **b)** bottom depth (i.e. bathymetry) in meters.

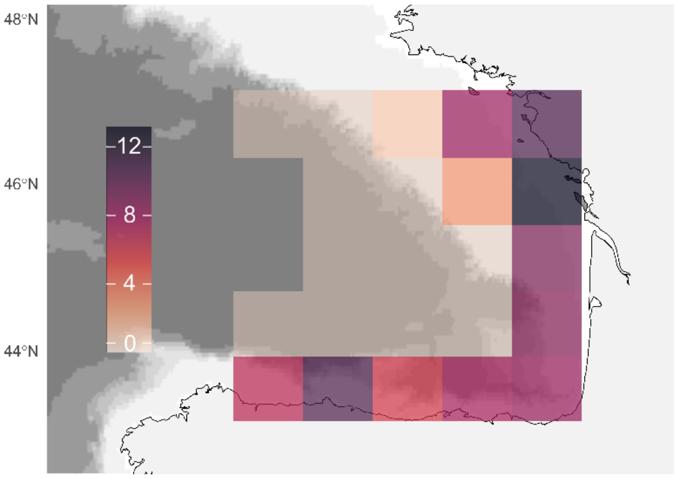


**Figure S8.** Depth-integrated mean of the acoustic model's posterior distribution of estimated anchovy biomass in logarithmic scale in the BoB. Negative values in logarithmic scale (corresponding to low biomass levels) were replaced by zero.

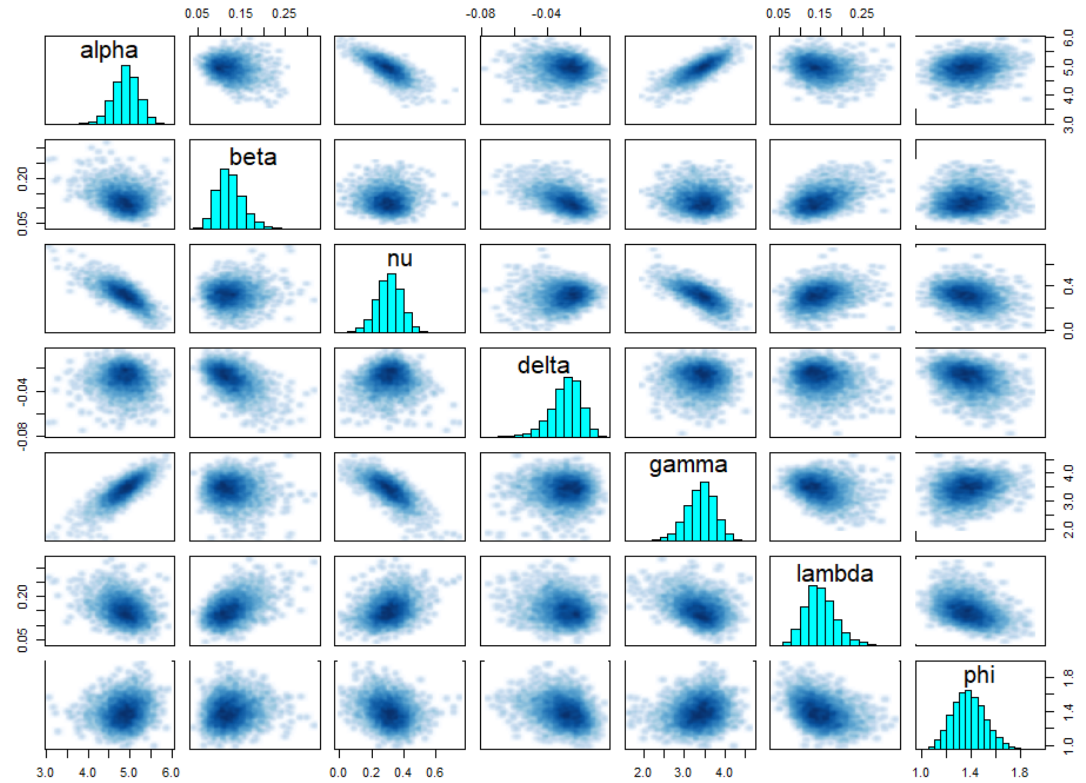


**Figure S5.** Pairs plot of the parameters estimated in the joint model.

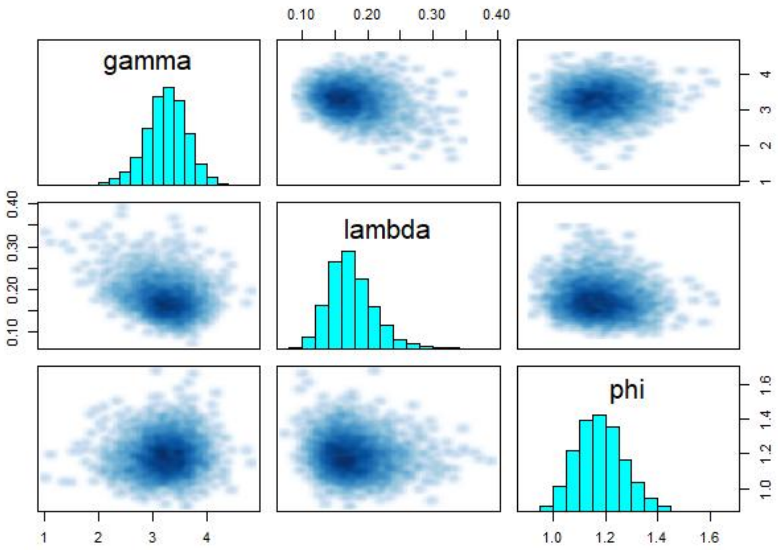


**Figure S8.** Pairs plot of the parameters estimated in the acoustic only model.

**Table S1.** Description of the data and parameters used in the joint model.

| | Description | Prior |
|---|---|---|
| **Data** | | |
| S | Number of cells in the grid | Not required |
| site_w | Index of the cells to which each acoustic value is assigned | Not required |
| site_y | Index of the cells to which each eDNA value is assigned | Not required |
| M | Total number of acoustic observations | Not required |
| N | Total number of eDNA observations | Not required |
| $\log_e W_i$ | Observed acoustic values in logarithmic scale in cell *i* | Not required |
| $\log_e Y_i$ | Observed eDNA values in logarithmic scale in cell *i* | Not required |
| **Parameter** | | |
| Theta ($\theta_i$) | Mean acoustic signal in cell *i* | N/A |
| Mu ($\mu_i$) | Mean eDNA concentration in cell *i* | N/A |
| Phi ($\varphi$) | Acoustic standard deviation | Gamma(0.1,0.1) |
| Sigma($\sigma_i$) | Genetic standard deviation in cell *i* | Gamma(0,5) |
| Gamma ($\gamma$) | Intercept of the linear relation between acoustic signal and biomass | Normal(-1,2) |
| Alpha ($\alpha$) | Intercept of the linear relation between eDNA concentration and biomass | Normal(0,5) |
| Nu ($v$) | Intercept of the linear relation between the standard deviation of eDNA predictions and biomass | Normal(-1,1) |
| Lambda($\lambda$) | Coefficient of $X_i$ for the observations of acoustic data | Normal(0,5) |
| Beta ($\beta$) | Coefficient of $X_i$ for the observations of eDNA data | Gamma(1,1) |
| Delta ($\delta$) | Coefficient of $X_i$ for the standard deviation of eDNA data | Normal(0,1) |
| **Other** | | |
| $X_i$ | Estimated (unobserved) biomass in logarithmic scale in the cell *i* | Normal(0,10) |
| **Subscripts** | | |
| i | Cell in grid (n = 23) | |

**Table S5.** Description of the data and parameters used in the acoustic model.

| | Description | Prior |
|---|---|---|
| **Data** | | |
| S | Number of cells in the grid | Not required |
| site_w | Index of the cells to which each acoustic value is assigned | Not required |
| M | Total number of acoustic observations | Not required |
| $\log_e W_i$ | Observed acoustic values in logarithmic scale in cell *i* | Not required |
| **Parameter** | | |
| Theta ($\theta_i$) | Mean acoustic signal in cell *i* | N/A |
| Phi (φ) | Acoustic standard deviation | Gamma(0.1,0.1) |
| Gamma (γ) | Intercept of the linear relation between acoustic signal and biomass | Normal(-1,2) |
| Lambda(λ) | Coefficient of $X_i$ for the observation of acoustic data | Normal(0,5) |
| **Other** | | |
| $X_i$ | Estimated (unobserved) biomass in logarithmic scale in the cell *i* | Normal(0,10) |
| **Subscripts** | | |
| i | Cell in grid (n = 23) | |

**Table S4.** Posterior means and 95% credible intervals of the parameters in the joint model.

| Parameter | Mean | 2.5% | 97.5% |
|---|---|---|---|
| alpha | 4.8904 | 4.2393 | 5.4590 |
| beta | 0.1242 | 0.0744 | 0.1962 |
| X[1] | -0.7622 | -8.3036 | 5.8692 |
| X[2] | 9.8263 | 3.5942 | 16.5462 |
| X[3] | -0.4742 | -7.8890 | 6.0788 |
| X[4] | 6.5514 | 1.2721 | 12.1949 |
| X[5] | 8.7348 | 3.0969 | 14.8896 |
| X[6] | 2.8530 | -5.6896 | 10.3561 |
| X[7] | -11.4678 | -20.8858 | -3.4345 |
| X[8] | -2.4045 | -11.0181 | 5.4416 |
| X[9] | -3.4065 | -11.6229 | 3.8725 |
| X[10] | 8.7478 | 2.0993 | 15.7009 |
| X[11] | -7.3485 | -20.7552 | 6.9266 |
| X[12] | -13.6514 | -23.6546 | -4.7406 |
| X[13] | 1.0316 | -5.9077 | 7.7366 |
| X[14] | 10.0550 | 3.3225 | 17.4380 |
| X[15] | -10.2219 | -19.3427 | -2.5307 |
| X[16] | -8.9424 | -16.5875 | -2.1257 |
| X[17] | 5.7459 | -1.7529 | 13.1251 |
| X[18] | 14.7284 | 6.5466 | 24.3147 |
| X[19] | -7.4259 | -20.7218 | 6.4228 |
| X[20] | -8.1828 | -16.8630 | -0.7136 |
| X[21] | 8.3681 | -0.5872 | 17.7077 |
| X[22] | 4.1162 | -3.4486 | 11.3304 |
| X[23] | 12.1164 | 3.9019 | 21.8591 |
| nu | 0.3086 | 0.1372 | 0.4662 |
| delta | -0.0291 | -0.0512 | -0.0140 |
| gamma | 3.4025 | 2.6174 | 4.0603 |
| lambda | 0.1491 | 0.0868 | 0.2323 |
| phi | 1.3790 | 1.1358 | 1.6586 |
| mu[1] | 4.8000 | 4.1119 | 5.3559 |
| mu[2] | 6.0660 | 5.5815 | 6.5302 |
| mu[3] | 4.8323 | 4.1298 | 5.3964 |
| mu[4] | 5.6760 | 5.3994 | 5.9387 |
| mu[5] | 5.9343 | 5.5818 | 6.2653 |
| mu[6] | 5.2633 | 4.3764 | 6.0836 |
| mu[7] | 3.5069 | 2.3148 | 4.4307 |
| mu[8] | 4.6160 | 3.6358 | 5.5256 |
| mu[9] | 4.4836 | 3.5274 | 5.2851 |
| mu[10] | 5.9438 | 5.3004 | 6.5604 |
| mu[11] | 3.9980 | 2.1157 | 5.8777 |
| mu[12] | 3.2529 | 1.9851 | 4.2814 |
| mu[13] | 5.0303 | 4.3219 | 5.7482 |
| mu[14] | 6.1029 | 5.4124 | 6.7793 |

| | | | |
|---|---|---|---|
| mu[15] | 3.6746 | 2.7149 | 4.4706 |
| mu[16] | 3.8393 | 3.2417 | 4.3782 |
| mu[17] | 5.5973 | 4.8247 | 6.4423 |
| mu[18] | 6.6925 | 5.7024 | 8.0500 |
| mu[19] | 3.9929 | 2.1625 | 5.7832 |
| mu[20] | 3.9038 | 2.8975 | 4.6813 |
| mu[21] | 5.9305 | 4.8741 | 7.1084 |
| mu[22] | 5.3808 | 4.6136 | 6.0296 |
| mu[23] | 6.3737 | 5.4394 | 7.7018 |
| theta[1] | 3.3441 | 2.6358 | 4.0973 |
| theta[2] | 4.8540 | 4.0988 | 5.6692 |
| theta[3] | 3.3809 | 2.6589 | 4.1096 |
| theta[4] | 4.3821 | 3.8171 | 5.0141 |
| theta[5] | 4.6953 | 4.0379 | 5.4201 |
| theta[6] | 3.8098 | 2.5907 | 4.7470 |
| theta[7] | 1.7676 | 0.6263 | 2.8220 |
| theta[8] | 3.0544 | 1.8329 | 4.0985 |
| theta[9] | 2.9276 | 1.9621 | 3.8167 |
| theta[10] | 4.6937 | 3.8859 | 5.5233 |
| theta[11] | 2.2824 | 0.0670 | 4.3997 |
| theta[12] | 1.4187 | -0.1743 | 2.8011 |
| theta[13] | 3.5586 | 2.6858 | 4.3445 |
| theta[14] | 4.8767 | 4.0309 | 5.7484 |
| theta[15] | 1.9365 | 0.7115 | 2.9659 |
| theta[16] | 2.1289 | 1.2210 | 2.9753 |
| theta[17] | 4.2383 | 3.3595 | 5.0988 |
| theta[18] | 5.5352 | 4.4948 | 6.5670 |
| theta[19] | 2.2713 | 0.0762 | 4.3134 |
| theta[20] | 2.2622 | 1.3719 | 3.0735 |
| theta[21] | 4.5873 | 3.4020 | 5.6229 |
| theta[22] | 4.0392 | 3.1576 | 4.9539 |
| theta[23] | 5.1561 | 4.1405 | 6.1782 |
| sigma[1] | 1.4082 | 1.2274 | 1.6961 |
| sigma[2] | 1.0399 | 0.8575 | 1.2334 |
| sigma[3] | 1.3928 | 1.2132 | 1.6524 |
| sigma[4] | 1.1394 | 1.0062 | 1.2899 |
| sigma[5] | 1.0721 | 0.9159 | 1.2413 |
| sigma[6] | 1.2605 | 0.9714 | 1.6084 |
| sigma[7] | 1.8998 | 1.5145 | 2.5062 |
| sigma[8] | 1.4607 | 1.1327 | 1.8509 |
| sigma[9] | 1.5063 | 1.2247 | 1.8764 |
| sigma[10] | 1.0702 | 0.8464 | 1.2938 |
| sigma[11] | 1.6950 | 1.0418 | 2.4857 |
| sigma[12] | 2.0126 | 1.5665 | 2.7014 |
| sigma[13] | 1.3242 | 1.0595 | 1.5913 |
| sigma[14] | 1.0318 | 0.7967 | 1.2726 |

| | | | |
|---|---|---|---|
| sigma[15] | 1.8280 | 1.4844 | 2.3291 |
| sigma[16] | 1.7623 | 1.4755 | 2.1521 |
| sigma[17] | 1.1608 | 0.8626 | 1.4237 |
| sigma[18] | 0.9062 | 0.5528 | 1.1970 |
| sigma[19] | 1.6997 | 1.0546 | 2.4936 |
| sigma[20] | 1.7320 | 1.4375 | 2.2060 |
| sigma[21] | 1.0790 | 0.7006 | 1.4180 |
| sigma[22] | 1.2224 | 1.0172 | 1.4561 |
| sigma[23] | 0.9741 | 0.6197 | 1.2590 |
| lp__ | -359.5986 | -371.4294 | -349.7483 |

**Table S5.** Posterior means and 95% credible intervals of the parameters in the acoustic model.

| Parameter | Mean | 2.5% | 97.5% |
|---|---|---|---|
| X[1] | 7.6277 | 0.8550 | 14.8138 |
| X[2] | 10.9444 | 4.1703 | 18.5438 |
| X[3] | 6.6582 | 0.0054 | 13.6938 |
| X[4] | 8.1475 | 1.4765 | 15.4002 |
| X[5] | 8.7707 | 1.9603 | 16.1813 |
| X[6] | -5.5214 | -14.7355 | 2.8795 |
| X[7] | -8.7085 | -17.5471 | -0.6612 |
| X[8] | -6.1235 | -15.0635 | 2.2190 |
| X[9] | -3.3678 | -10.6031 | 3.4328 |
| X[10] | 9.3128 | 2.6078 | 16.4853 |
| X[11] | -12.7143 | -24.8932 | -1.2422 |
| X[12] | -15.0157 | -25.6639 | -5.2917 |
| X[13] | -1.8373 | -8.9085 | 4.6791 |
| X[14] | 9.5605 | 2.9182 | 17.0549 |
| X[15] | -8.3680 | -18.5808 | 1.0656 |
| X[16] | -8.5426 | -16.9393 | -1.1197 |
| X[17] | 3.4038 | -3.3927 | 10.1090 |
| X[18] | 13.6236 | 6.4104 | 21.6244 |
| X[19] | -12.6749 | -24.5959 | -0.9788 |
| X[20] | -0.4773 | -8.3800 | 7.0406 |
| X[21] | 1.7287 | -5.4678 | 8.7173 |
| X[22] | 8.5818 | 1.4780 | 16.1573 |
| X[23] | 11.1207 | 4.2327 | 18.8819 |
| gamma | 3.2567 | 2.4176 | 3.9878 |
| lambda | 0.1753 | 0.1179 | 0.2590 |
| phi | 1.1808 | 1.0143 | 1.3808 |
| theta[1] | 4.5665 | 3.6502 | 5.4682 |
| theta[2] | 5.1294 | 4.2074 | 6.0320 |
| theta[3] | 4.4017 | 3.4845 | 5.3145 |
| theta[4] | 4.6559 | 3.7557 | 5.5573 |
| theta[5] | 4.7605 | 3.8586 | 5.6833 |
| theta[6] | 2.3310 | 1.0636 | 3.5890 |
| theta[7] | 1.7910 | 0.6982 | 2.9243 |
| theta[8] | 2.2270 | 0.9682 | 3.4668 |
| theta[9] | 2.7030 | 1.7629 | 3.6414 |
| theta[10] | 4.8537 | 3.9341 | 5.7476 |
| theta[11] | 1.0687 | -0.9156 | 3.0953 |
| theta[12] | 0.6979 | -0.8266 | 2.2605 |
| theta[13] | 2.9619 | 2.0674 | 3.8798 |
| theta[14] | 4.8954 | 3.9863 | 5.7939 |

| | | | |
|---|---|---|---|
| theta[15] | 1.8379 | 0.3161 | 3.3615 |
| theta[16] | 1.8215 | 0.8411 | 2.8304 |
| theta[17] | 3.8503 | 2.9160 | 4.7812 |
| theta[18] | 5.5835 | 4.6695 | 6.4768 |
| theta[19] | 1.0764 | -0.9327 | 3.0841 |
| theta[20] | 3.1906 | 2.0957 | 4.2761 |
| theta[21] | 3.5678 | 2.5546 | 4.5729 |
| theta[22] | 4.7295 | 3.7400 | 5.7141 |
| theta[23] | 5.1585 | 4.2500 | 6.0642 |
| lp__ | -85.9889 | -97.6679 | -75.9438 |
